# Transcranial ultrasound stimulation of the medial frontal cortex transiently improves performance by modulating conflict monitoring dynamics

**DOI:** 10.64898/2026.08.06.743260

**Authors:** Camila S. Agostino, Hans Kirschner, Daniel Janko, Eleonora Carpino, Lennart Verhagen, Markus Ullsperger

## Abstract

Conflict monitoring and error processing are fundamental mechanisms underlying cognitive control and adaptive behavior and have been consistently associated with increased activity in the anterior midcingulate cortex (aMCC). Here, we used transcranial ultrasound stimulation (TUS), an emerging technique that enables non-invasive, deep, and focal neuromodulation, and EEG to investigate the causal role of the aMCC in cognitive control. Our findings demonstrate that TUS of aMCC improved performance, modulated the relationship between conflict monitoring and the stimulus-locked N2, and strengthened the suppression of distracting flankers as revealed by drift-diffusion model analyses. This suggests that TUS of aMCC enhances proactive control. Interestingly, TUS did not affect the error-related negativity as a measure of error monitoring. Finally, the TUS effects were observed during early compared with later task blocks corroborating previous findings which suggested that TUS effects are temporally dynamic and characterized by a limited post-stimulation window.

**HIGHLIGHTS:**

- aMCC-TUS and PCC-TUS both increase behavioral accuracy during the early stages of task performance.
- aMCC-TUS transiently reduces the effect of incongruence on the stimulus-locked N2 suggesting reduced response conflict.
- Exploratory DDM analyses suggest that aMCC-TUS improves suppression of conflict-inducing distractors.
- aMCC thus selectively enhances proactive control, thereby reducing response conflict.
- TUS effects show a transient temporal profile, peaking 17–27 minutes after stimulation and declining after ∼37 minutes.

## INTRODUCTION

Cognitive control, specifically response conflict monitoring and error processing, has been extensively researched throughout recent decades. And although the vast majority of the studies are correlational, few causal findings were established either via non-invasive brain stimulation (NIBS) techniques or lesion studies, which have provided valuable insights into brain function, helping to clarify the contribution of specific regions and their interactions within broader networks. However, interpretations based solely on lesion evidence must be made with caution, as neural plasticity and compensatory reorganization in patients can obscure causal relationships. Commonly used NIBS techniques such as transcranial magnetic stimulation (TMS) and transcranial direct current stimulation (tDCS) offer alternative causal approaches, but their limited spatial specificity and restricted ability to target deeper structures constrain the precision with which individual regions can be investigated. In this context, transcranial ultrasound stimulation (TUS) has emerged as a particularly promising tool, enabling high-resolution modulation of both cortical and subcortical areas and offering a unique opportunity to probe their causal roles in functions such as error processing and conflict monitoring.

Processing errors and monitoring response conflict are essential components of cognitive function, allowing individuals to detect and correct erroneous responses to maintain goal-directed behaviour (Dehaene et al., 1994; Carter et al., 1998, Ullsperger & von Cramon, 2001; Ridderinkhof et al., 2004; Debener et al., 2005; Fu et al., 2022). Electrophysiological studies have identified event-related potentials (ERP) associated with errors, the error-related negativity (ERN) or error negativity (Ne) (Falkenstein et al., 1991; Gehring et al., 1993). Observed at central electrodes, the ERN deflects around 0 and 100 ms after the response and has been associated with detecting discrepancies between intended and executed actions, prompting necessary behavioural adjustments (Taylor et al., 2007; Ullsperger & von Cramon, 2004; Vilà-Balló et al., 2014). Following ERN, an error positivity (Pe) component is observed in frontocentral regions and later, in centroparietal areas (Falkenstein et al., 1991) and it is usually associated with conscious error perception (Ullsperger et al., 2014; Ullsperger, 2024). Response conflict evolving during response selection is reflected in an amplitude increase of the stimulus-locked frontocentral N2 (van Veen & Carter, 2002a; Folstein & van Petten, 2008; Ullsperger et al., 2014a; Ullsperger et al 2014b), which deflects around 200 and 350 ms, or longer depending on the task complexity (Brass et al., 2005; Swick & Turken, 2002; West, 2003). Neuroimaging studies revealed multiple regions and networks, each contributing to different aspects of error detection (Ullsperger & von Cramon, 2001; Ullsperger et al., 2014a) and response conflict (Cieslik et al., 2024). One of the most consistently implicated regions is the anterior midcingulate cortex (aMCC), also part of the salience network that responds to behaviourally significant events (Neta et al., 2015; Holroyd & Coles, 2002; Ham et al., 2013, Kirschner & Ullsperger, 2025). Source localization, simultaneous EEG/fMRI recordings and invasive recordings suggest that both the ERN and the frontocentral N2 are generated at least in part in the aMCC (Dehaene et al., 1994; Gruendler et al., 2011; Wessel et al., 2012; Fu et al., 2019; Fu et al., 2022). Changes in neural correlates of performance monitoring and behavioral maladaptation have been linked to various disorders, such as schizophrenia (Kirschner & Klein, 2022), ADHD (Cai et al., 2021) and Obsessive Compulsive Disorder (Endrass & Ullsperger, 2014). Therefore, understanding the mechanisms related to performance monitoring, especially error processing, is of utmost importance to promote more precise treatment for such disorders.

Lesion studies open a door to investigate this causal relation, but focal lesions in the aMCC are rare and evidence is sparse. One study reported a single case of a patient with a lesion on the rostral to mid-aMCC, who presented smaller ERN compared to healthy controls, while the conflict-related modulation of the N2 was unaffected (Swick & Turken, 2002). Similarly, another study revealed a direct connection between ventrolateral anterior thalamus and aMCC by showing that patients with thalamic lesions also presented smaller ERN (Seifert et al., 2011). In healthy brains, on the other hand, few studies attempted to modulate the aMCC, as this region is relatively hard to reach for most NIBS techniques. For instance, in a TMS study using a double cone coil, Hayward and colleagues (2004) targeted the aMCC in order to investigate behavioural changes in a counting Stroop task. A significant increase in reaction time was observed in incongruent trials after TMS was applied over the motor cortex (control region), while no difference in reaction time during congruent and incongruent trials was observed after stimulation of different portions of the aMCC, suggesting that TMS over this region abolished the interference effect in Stroop task. In a transcranial electrical stimulation study, Mattavelli et al. (2022) applied high-definition tDCS to the aMCC to examine performance monitoring. Results indicated that cathodal aMCC stimulation significantly modulated performance in a Flanker Task, by decreasing RT in incongruent trials compared to anodal stimulation, and increasing RT in congruent trials compared to sham, suggesting an increasing control over the Flanker conflict effect. Although TMS with specific coils and high-definition tDCS can be used for reaching aMCC, these techniques are not ideal, as they lack spatial specificity, meaning that the stimulation field can reach other undesired regions.

Transcranial ultrasound stimulation (TUS), on the other hand, has been explored in humans in the last years (Darmani et al., 2022; Murphy & Fouragnan, 2024; Murphy et al., 2025) and has showed promising outcomes when targeting the cingulate cortex. Yaakub and colleagues (2023) investigated functional connectivity in resting state after stimulating the aMCC and posterior cingulate cortex (PCC) with the offline 5Hz rTUS protocol (Zeng et al. 2021). The authors observed an increase in connectivity in aMCC after stimulation of this region, but also TUS of PCC, compared to sham. However, connectivity in PCC was observed only after stimulation of the same region and not after stimulation of aMCC. Connectivity results also revealed an increase in the salience network after stimulation of the aMCC and an increase in connectivity in the default mode network after stimulation of the PCC. A reduction in GABA in the PCC was seen after TUS of this region, providing one more evidence of the effectiveness of TUS application.

In the current study, we used TUS to target the aMCC, followed by a modified version of Eriksen’s Flanker task and continuous EEG recording, to examine its causal relationship to error commission and conflict monitoring. As control conditions, PCC was targeted as an active control region, and a sham protocol was delivered to the aMCC. We hypothesised that TUS of aMCC would modulate performance, as well as the electrophysiological markers, such as the ERN or Pe, differently compared to TUS of PCC and sham. Although there is still limited evidence regarding the directionality of TUS effects for the protocol used here, we expected a facilitatory effect, meaning an enhancement of conflict-control processes. Specifically, we employed a 5-Hz stimulation protocol that has previously been suggested to lead to facilitatory effects when applied to our regions of interest (Yaakub et al., 2023).

## RESULTS

Nineteen participants (mean age = 26 years, SD = 2.57; six women; one left-handed man) took part in a five-day study (Fig. 1.A) comprising one fMRI session, three TUS+EEG sessions, and one EEG session. Across all sessions, participants performed an arrow-based version of the Eriksen Flanker Task adapted from Fischer et al. (2018). On each trial, four flanker arrows were presented prior to the appearance of a five-arrow array. Participants were instructed to respond as quickly and accurately as possible to the orientation of the central arrow, which could be either congruent (same orientation) or incongruent (opposite orientation) relative to the flankers. During the MRI session, structural images were acquired and used to carry out thermal and acoustic ultrasound simulations. Functional images were used to determine individual stimulation targets. During the TUS sessions, participants received a 5-Hz repetitive stimulation protocol for 120 seconds targeting either the aMCC or PCC, as well as a sham stimulation over the aMCC lasting 2 seconds, prior to the EEG and behavioural assessments. The aMCC target was functionally localized using the contrast incongruent incorrect > congruent correct trials (Fig. 1B) bilaterally for each subject, whereas PCC masks were defined based on the absence of activation in the same contrast within the posterior cingulate cortex region (Fig. 1C).

**Figure 1.**
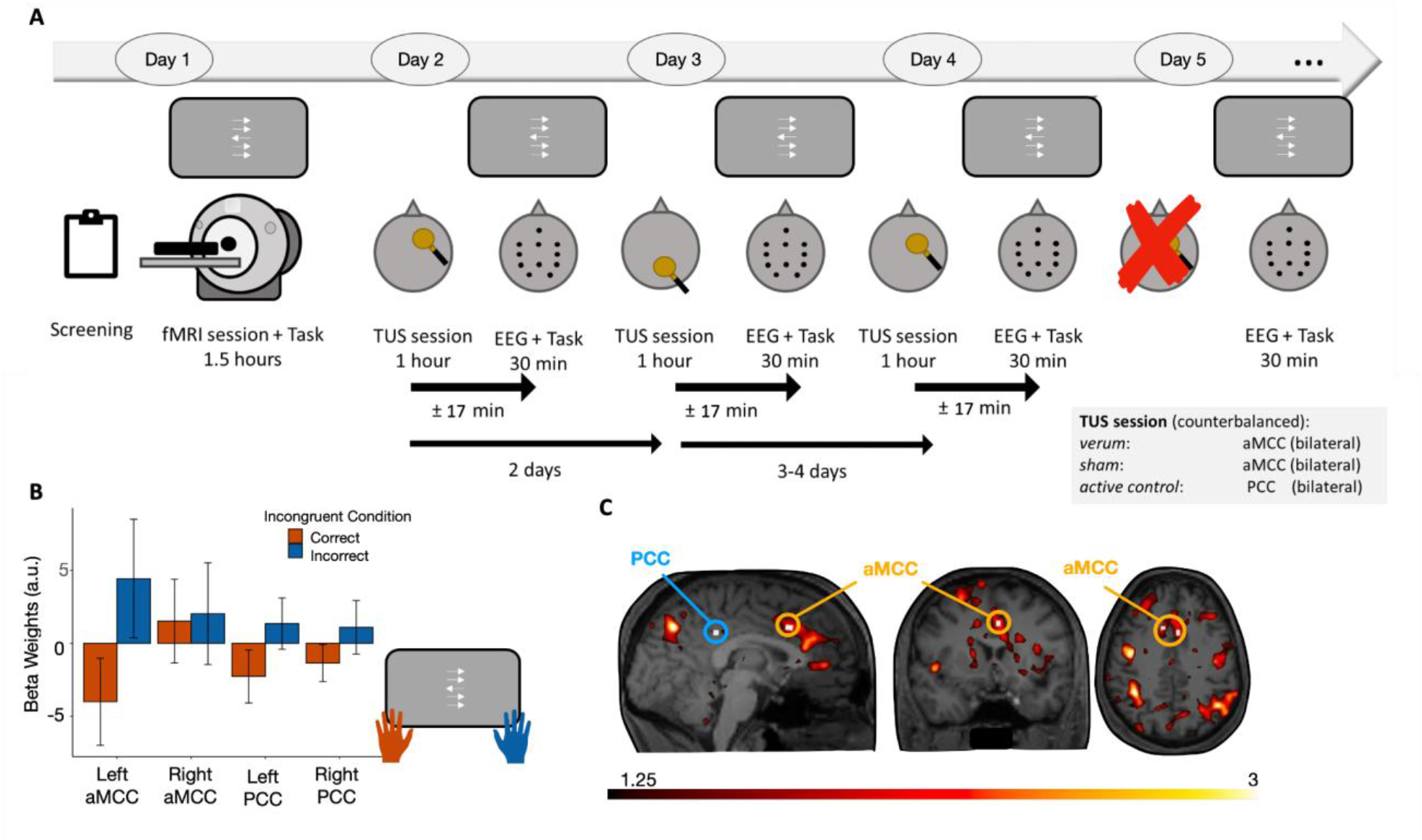
Experimental Design and fMRI results: **A)** Participants took part in one fMRI session in which structural and functional data were acquired while they performed the Eriksen Flanker task. On different days, participants underwent three TUS sessions, in which they received the 5Hz rTUS protocol for 120s targeting the aMCC or the PCC, as well as a 2 second stimulation of the aMCC as our “sham” protocol. Following the stimulation, a reduced-EEG set-up was prepared and participants performed again the Flanker task. The time between the end of the stimulation and beginning of EEG measurement lasted on average 17 min (SD=±1.51) and the interval between stimulation sessions varied between 2 and 4 days. **B)** Activation pattern of the beta weights from incorrect and correct incongruent trials averaged across participants are shown. **C)** Anatomical image of an exemplary subject shows the overlay of contrast cluster of incorrect > correct incongruent trials. The white squares circled by the orange and blue circles highlight the aMCC and PCC masks, respectively, which were created based on functional and anatomical MRI data. The aMCC masks were selected from the subjects’ regions local maxima of the contrast incorrect incongruent > correct incongruent, while the PCC masks were created from the region with minimum to none activation in the same contrast.

Before each individual TUS session, acoustic simulations were conducted to estimate stimulation parameters. Across stimulation conditions, the calculated average spatial-peak pulse-average intensity (Isppa; indicated as Ipa in Table 1 of the Methods section) at the target site was 7.56 W/cm² (Fig. 2A.1). In addition, dose calculations (table 1) resulted in negligible energy absorption for the sham condition (mean = 1.1, SD = +-0.2) relative to the active aMCC (87.4, +-4.8) and PCC (75.6, +-6.8) stimulation conditions (Fig. 2A.2). All simulations reached the regions of interest as indicated by the exemplary subject in figure 2.A.3. The individual masks were normalized across subjects and overlapped in order to verify if all participants shared voxels around the regions of interest (Fig. 2.A.4).

**Figure 2.**
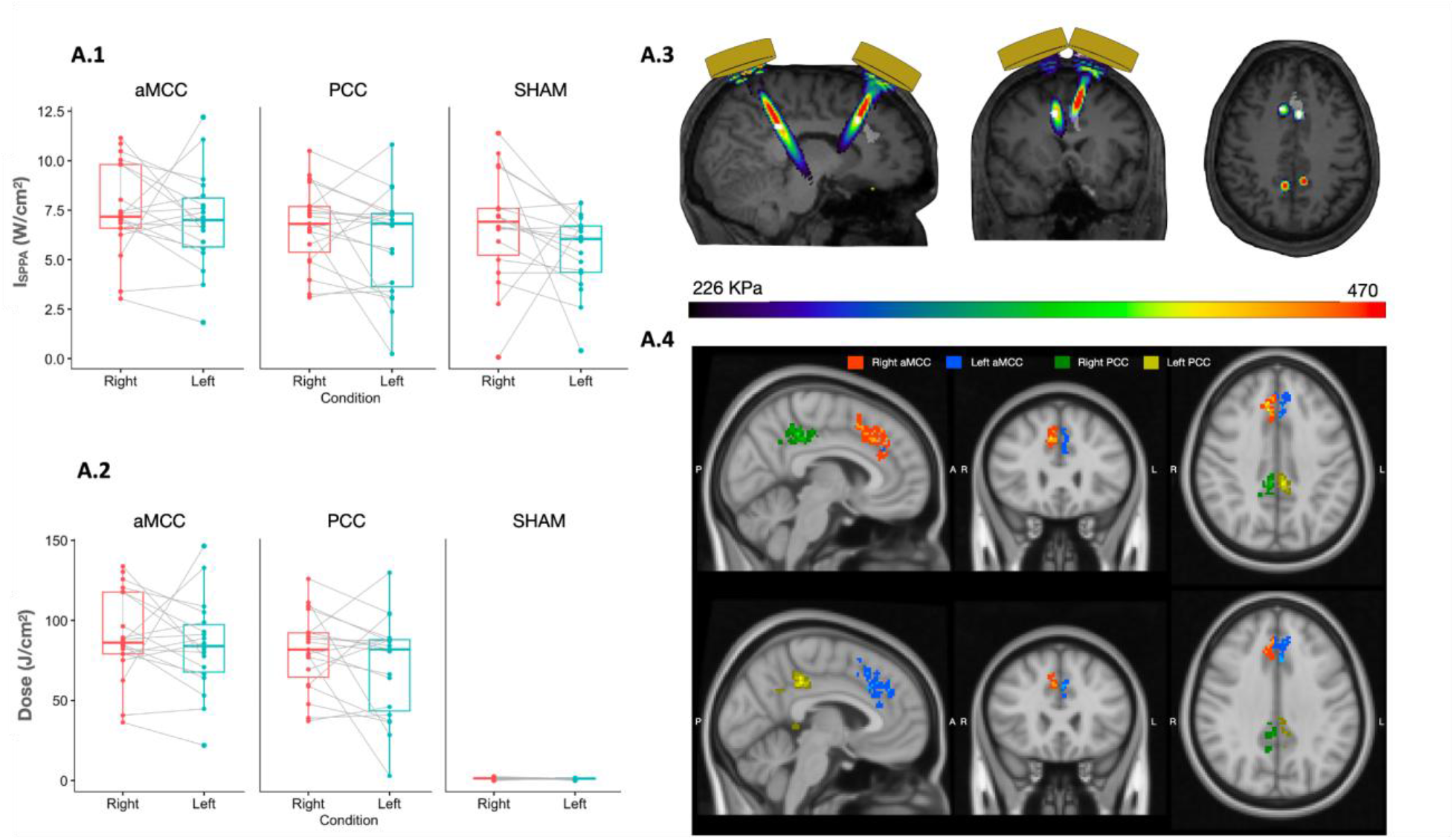
Simulation Information: **A.1)** ISPPA of left and right aMCC, PCC and sham are depicted. Note that we present the ISPPA of the sham condition because here the stimulation protocol was delivered for 2 seconds, so the participants could experience the auditory confound similarly to the verum stimulations. **A.2)** The dose was calculated based on Nandi et al. (2025) and shows that the exposure was near zero during sham condition. **A.3)** Simulation targets of bilateral aMCC and PCC are shown on an exemplary subject. **A.4)** Overlay of normalized individual bilateral aMCC (red = right; blue = left) and PCC (green = right; yellow = left) masks over normalized brain. The overlays show that participants received stimulation over the pMFC and PCC regions, but also a portion of Precuneus. The difference in contrast represents the number of subjects which shared the same voxels (the brighter the colour, the higher the number of participants sharing that ROI location).

**Figure 3.**
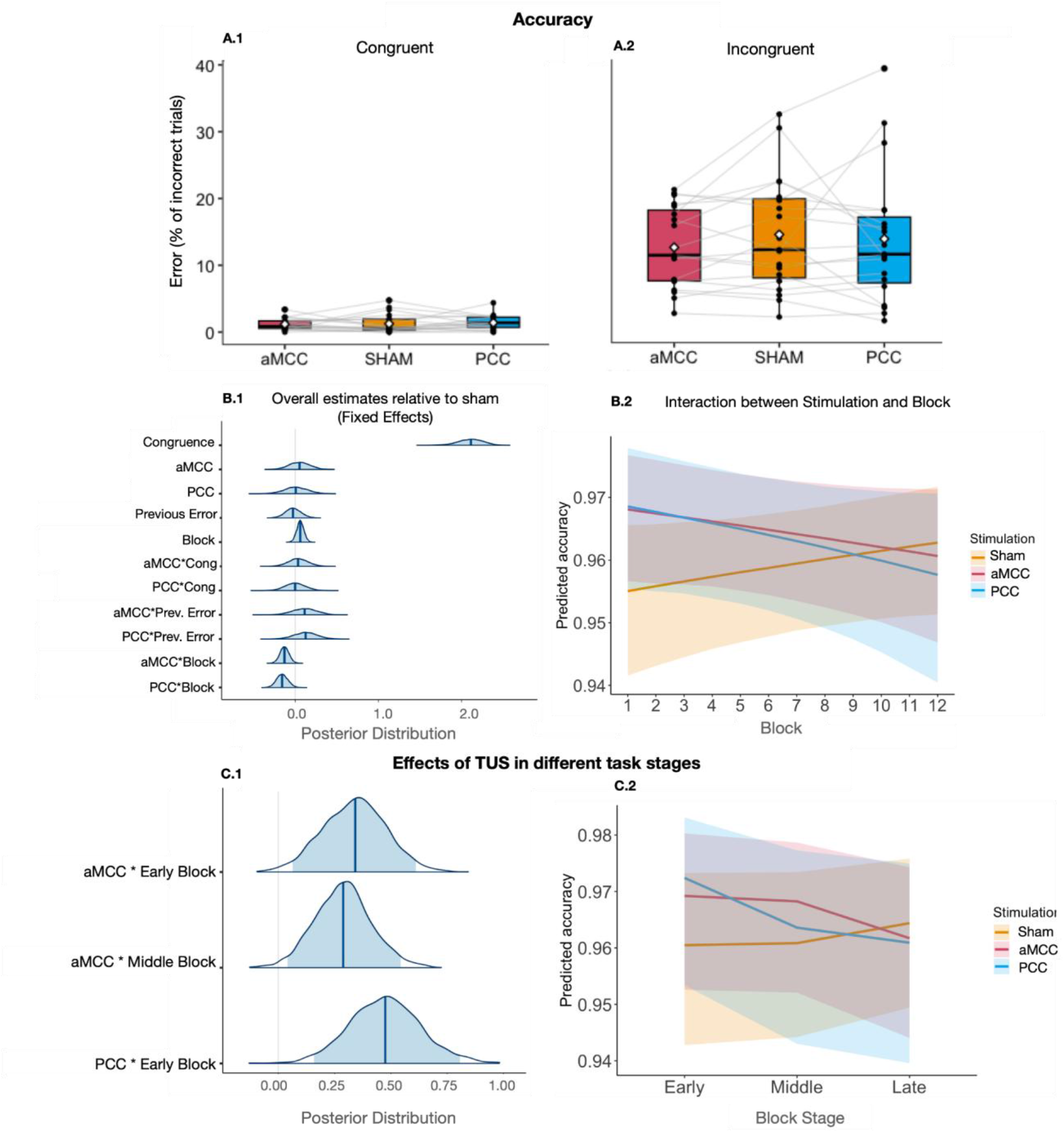
Accuracy Model Results: **A.1)** Error rates in congruent trials and **A.2)** incongruent trials. **B.1)** All valid trials were fitted in the model which revealed a strong effect of congruence and a credible interaction between aMCC-TUS and Blocks, as well as PCC-TUS and blocks. **B.2)** After aMCC-TUS and PCC-TUS, relative to sham, predicted accuracy was higher in the beginning of the task compared to the end of the task. **C.1)** The twelve blocks were divided in three windows: earlier, middle and later blocks (1 to 4, 5 to 8 and 9 to 12, respectively) and fitted into a model, which revealed differences between earlier and middle blocks relative to later blocks, after aMCC-TUS. After PCC-TUS, accuracy was different in earlier blocks compared to later ones. **C.2)** Predicted accuracy was higher in earlier blocks compared to later blocks after aMCC-TUS and PCC-TUS compared to sham.

**Figure 4.**
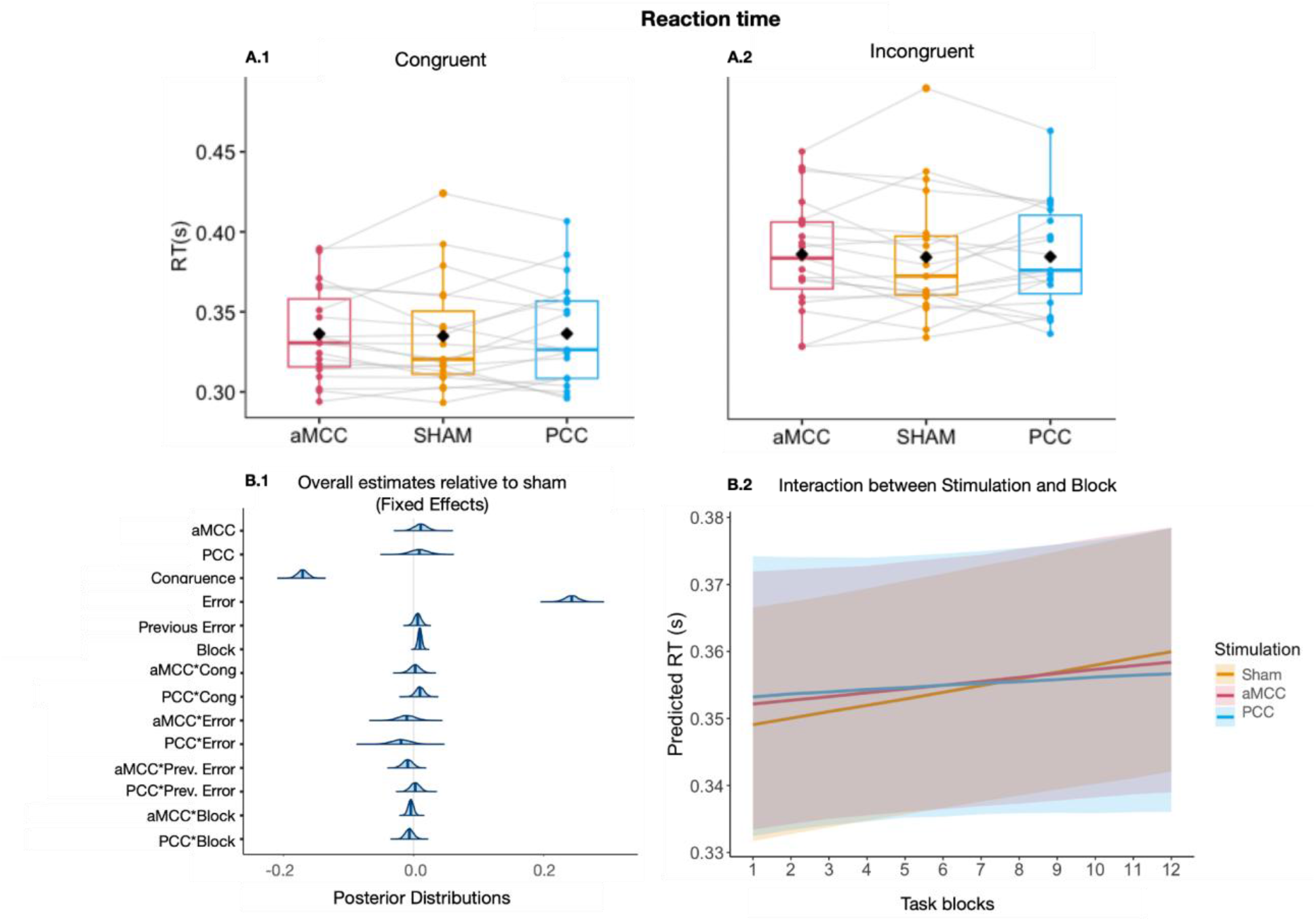
Reaction Time Model Results. **A.1)** Reaction times for congruent trials were shorter than for incongruent trials (**A.2**). **B.1)** All valid trials were included in the reaction time model, which revealed a strong effect of congruence and error, indicating that participants responded faster to congruent trials and slower to correct trials. **B.2)** Reaction time was kept relatively constant across stimulation conditions over time.

**Table 1.** Table with acoustic and thermal parameters following the ITRUSST safety recommendations. . Ptp target represents the pressure in the location of the stimulation target with potential offset. Temperature below the skull was always below 38℃, as we assumed a baseline to 37℃, thermal dose (CEM43) was always below 0.002 in the brain and MItc stands for transcranial Mechanical Index.

|  | aMCC |  | PCC |  |
| --- | --- | --- | --- | --- |
|  | Right | Left | Right | Left |
| **Ptp target (kPA)** | 487.9 ($\pm 62.33$) | 452.83 ($\pm 75.40$) | 468.44($\pm 77.19$) | 476.16 ($\pm 63.26$) |
| **lpa (W/cm2)** | 8.05 (1.96) | 7.01 ($\pm 2.34$) | 7.5 ($\pm 2.28$) | 7.68 ($\pm 1.97$) |
| **Ispta (W/cm2)** | 1.04 ($\pm 0.17$) | 1.20 ($\pm 0.52$) | 1.36 ($\pm 0.24$) | 1.39 ($\pm 0.39$) |
| **Isppa (W/cm2)** | 11.95 ($\pm 2.04$) | 13.80 ($\pm 6.00$) | 15.63 ($\pm 2.96$) | 15.82 ($\pm 4.06$) |
| **ROI Peak Pressure (kPA)** | 470.85 ($\pm 77.24$) | 450.50 ($\pm 86.11$) | 443.23 ($\pm 71.13$) | 406.15 ($\pm 113.20$) |
| **ROI Mean Pressure - 6db (kPA)** | 342.24 ($\pm 63.35$) | 325.02 ($\pm 68.40$) | 350.84 ($\pm 67.01$) | 326.50 ($\pm 105.4$) |
| **-6 db focal volume (mm3)** | 431.17 ($\pm 303.92$) | 505.17 ($\pm 413.31$) | 126.52 ($\pm 102.43$) | 95.25 ($\pm 36.11$) |
| **Mltc** | 1.11 ($\pm 0.11$) | 1.12 ($\pm 0.14$) | 1.27 ($\pm 0.14$) | 1.23 ($\pm 0.18$) |
| **Focal distance (mm)** | 46.4 ($\pm 3.14$) | 46.4 ($\pm 3.14$) | 57.36 ($\pm 3.43$) | 57.36 ($\pm 3.43$) |
| **Dose (J/cm2)** | 90.8 ($\pm 27.2$) | 84.0 ($\pm 29.7$) | 80.5 ($\pm 24.4$) | 70.8 ($\pm 31.5$) |

Performance was assessed via reaction time and accuracy measures, which were analysed using Bayesian linear mixed-effects models. We included congruence, previous error, block (plus accuracy for the reaction time model), and their interactions with stimulation conditions (sham, aMCC, PCC) as fixed effects and allowed them to vary on the subject level (random effect of subject). This allowed us to assess the impact of conflict processing, post-error slowing, and task progression on the dependent variable.

### TUS dynamically modulated behaviour over tim e

Descriptive results demonstrated a clear difference in error commission between congruent (Fig. 3.A.1) and incongruent trials (Fig. 3.A.2). The accuracy model (see Eq. 2.1 in the Methods section; Fig. 3.B.1) did not reveal a general credible main effect of stimulation, with neither aMCC-TUS (β = 0.05, 95% CrI [−0.19, 0.29]) nor PCC-TUS (β = 0.01, 95% CrI [−0.27, 0.29]) differing from sham. However, credible stimulation-block interactions showed that accuracy varied over the course of the task (Fig. 3.B.2), after aMCC-TUS (β = −0.13, 95% CrI [−0.24, −0.02]) and PCC-TUS (β = −0.16, 95% CrI [−0.29, −0.02]), reversing the direction of the effect block had in the sham condition. A robust effect of congruence under sham stimulation (β = 2.11, 95% CrI [1.78, 2.38]) was observed, indicating higher accuracy for congruent relative to incongruent trials. Other effects on task dynamics, such as post-error adjustments were not observed as previous errors did not reliably influence current-trial accuracy (β = −0.02, 95% CrI [−0.21, 0.16]). TUS also did not influence other factors as no credible interactions were seen between stimulation and congruence (aMCC-TUS × congruence: β = 0.04, 95% CrI [−0.20, 0.27]; PCC-TUS × congruence: β = 0.00, 95% CrI [−0.24, 0.24]) or between stimulation and previous error (aMCC-TUS × previous error: β = 0.11, 95% CrI [−0.18, 0.39]; PCC-TUS × previous error: β = 0.13, 95% CrI [−0.15, 0.40]). These results confirm a robust conflict effect while suggesting that TUS modulated performance dynamics over time, by showing that stimulation was associated with improved accuracy early in the task, an effect that diminished across later blocks. This partially aligns with our hypothesis that aMCC-TUS modulates behavioural performance. However, PCC-TUS seems to also have affected accuracy in the beginning of the task.

To further characterise the temporal profile of TUS effects, we fitted an additional model (Eq. 2.3 in the Methods section) in which task blocks were grouped into three time windows: early (blocks 1–4; ∼17 min from the end of TUS to the onset of the first block), middle (blocks 5–8; ∼27 min post-TUS), and late (blocks 9–12; ∼37 min post-TUS). The model revealed higher accuracy in the early relative to late window for both aMCC-TUS (β = 0.34, 95% CrI [0.06, 0.61]) and PCC-TUS (β = 0.47, 95% CrI [0.16, 0.84]) compared to sham (Fig. 3.C.1). A similar, albeit smaller, effect was observed in the middle window for aMCC-TUS relative to sham (β = 0.29, 95% CrI [0.04, 0.54]). Figure 3.C.2 illustrates this shift in the accuracy over blocks. These findings indicate a time-limited effect of TUS on behaviour, with performance benefits that are strongest around 17 to 27 min after stimulation and diminish after 37 minutes.

### TUS did not affect reaction time

Congruent and incongruent trials descriptively differed in reaction time (RT) across participants (Fig 4.A1-2). A robust effect of response accuracy was observed in the reaction time model (see Eq. 2.2 in the Methods section, Fig. 4.B.1), under sham stimulation (reference), with slower responses on correct compared to erroneous trials (β = 0.24, 95% CrI [0.22, 0.26]), as well as a clear effect of congruence, with faster responses on congruent compared to incongruent trials (β = −0.17, 95% CrI [−0.19, −0.15]). A small but credible slowing across blocks was evident under sham (β = 0.01, 95% CrI [0.00, 0.02]). In contrast to the accuracy model, this was not further modulated by either of the stimulation conditions as no credible interaction between stimulation and block progression was observed (aMCC-TUS × block: β = 0.00, 95% CrI [−0.01, 0.00]; PCC- TUS × block: β = −0.01, 95% CrI [−0.02, 0.01]). Descriptively (Fig. 4.B.2), reaction times under sham increased towards the end of the task, whereas responses following stimulation remained comparatively stable; however, these differences were minimal and not credible. Additionally, there was no evidence for a main effect of stimulation (aMCC-TUS: β = 0.01, 95% CrI [−0.01, 0.03]; PCC-TUS: β = 0.01, 95% CrI [−0.02, 0.04]) or previous error (β = 0.01, 95% CrI [−0.00, 0.02]). Similarly, no credible interactions were observed between stimulation and congruence (aMCC-TUS × congruence: β = 0.00, 95% CrI [−0.01, 0.02]; PCC-TUS × congruence: β = 0.01, 95% CrI [−0.00, 0.02]), response accuracy (aMCC-TUS × error: β = −0.01, 95% CrI [−0.04, 0.02]; PCC-TUS × error: β = −0.02, 95% CrI [−0.05, 0.01]), or previous error (aMCC-TUS × previous error: β = −0.01, 95% CrI [−0.02, 0.01]; PCC-TUS × previous error: β = 0.00, 95% CrI [−0.01, 0.02]). Finally, the model confirmed that incongruent trials elicited slower responses and that error trials were associated with faster responses relative to correct trials. Crucially, stimulation did not influence overall response speed, conflict processing, or post-error slowing, not supporting our initial hypothesis.

### Performance scales with TUS intensity

To explore the nature of inter-individual variability observed in the data, we conducted correlation analyses between the subject-level estimates of different model parameters and individual delivered ultrasound intensity (Isppa, Fig. 2.A.1). Results revealed a dose-dependent effect particularly after PCC-TUS. Model-estimated inter-individual variability in difference in RT and accuracy between PCC-TUS and sham was positively correlated with Isppa (Fig. 5.A - accuracy: r = 0.65, p = 0.003; Fig. 5.B - RT: r = 0.46, p = 0.046), suggesting that intensity may explain individual differences in the effect of PCC stimulation.

**Figure 5.**
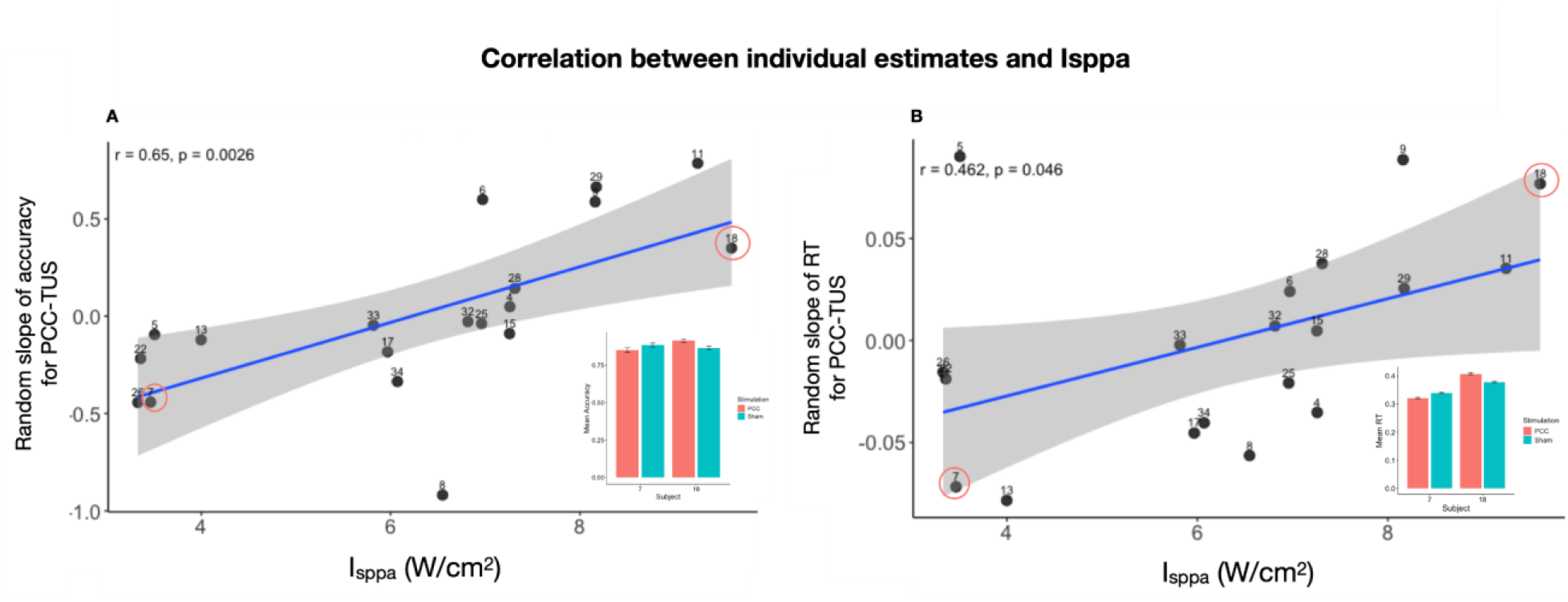
Behaviour and ISSPA Correlation: **A**) Accuracy positively correlated with Isppa after PCC- TUS. The red and green bars represent the mean accuracy after PCC-TUS and sham, respectively, for the subject which received the weakest and the strongest intensity (subjects 7 and 18). **B)** Reaction time positively correlated with Isppa, indicating that higher intensities led to slower RTs after PCC-TUS. The red and green bars represent the mean RT after PCC-TUS and sham, respectively, for the subject which received the weakest and the strongest intensity (subjects 7 and 18).

### TUS selectively affects conflict-related but not error-related neural processing

#### ERN and error commission relation was not modulated by aMCC-TUS

A strong ERN component was elicited across all sessions (Fig. 6.A). In order to investigate the difference in the ERN profile across participants and sessions, we conducted single-trial regression analysis which revealed positive estimates (Fig. 6.B) across all stimulation conditions, indicating a robust ERN effect, with more negative amplitudes for error relative to correct trials. Bayesian mixed-effects modeling revealed no credible difference between sham and aMCC-TUS condition (β=-0.09, 95% CrI [-0.76, 0.59]; or PCC-TUS (β=0.67, CrI 95%[-0.09, 1.43]). Pairwise comparison revealed marginally a credible difference between aMCC-TUS and PCC-TUS (β=- 0.7583 95% CrI [−1.553 0.0501]; Fig. 6.C).

**Figure 6.**
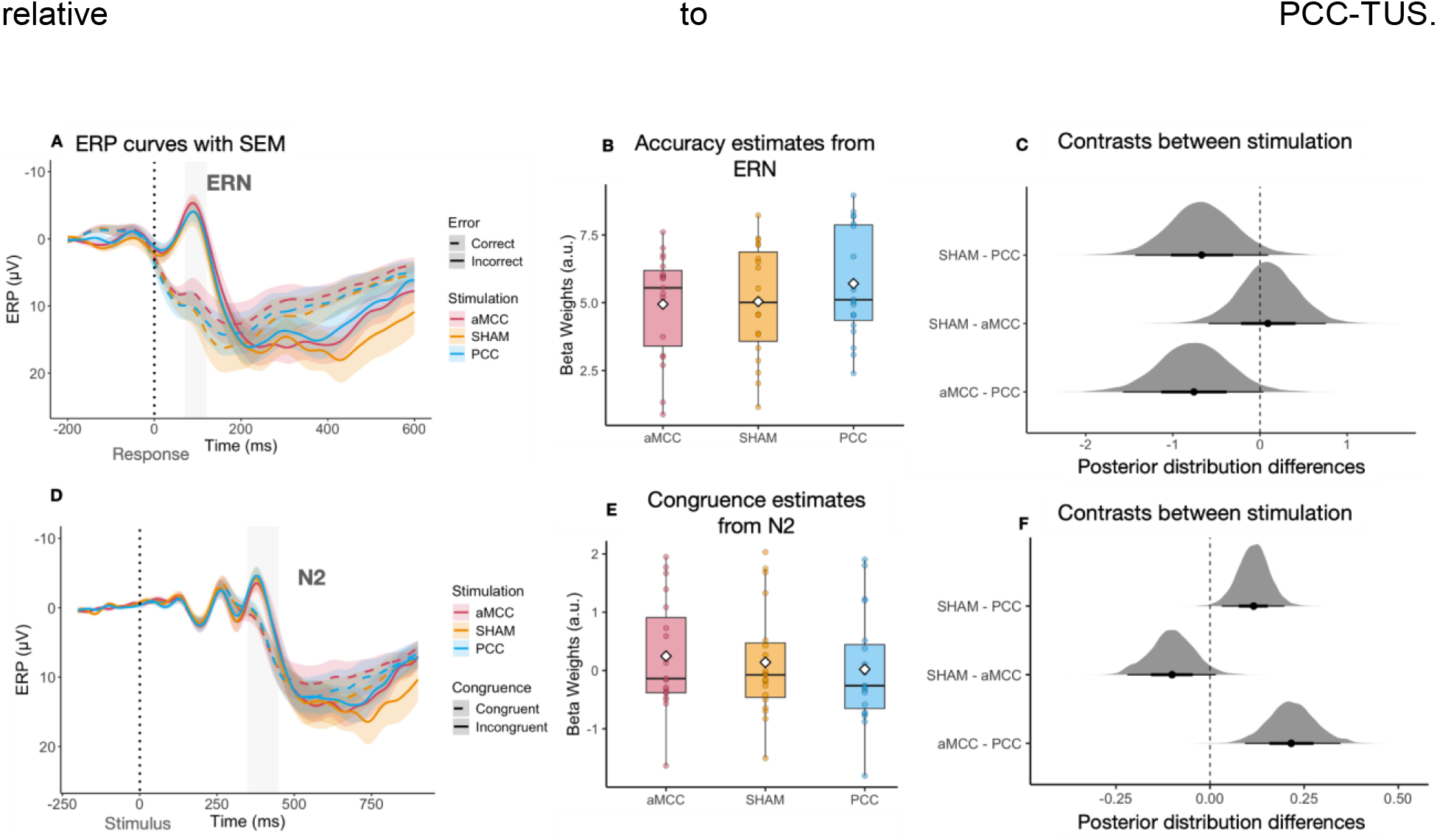
ERP Model Results: **A)** The curves represent the event-related potential (ERP) with SEM after aMCC-TUS (red), PCC-TUS (blue) and sham (yellow) conditions for correct (dashed line) and incorrect (full line) trials. An ERN was elicited after errors. **B)** The boxplot shows the betas of single-trial regression analysis were averaged across the analysis interval (70-120 ms; light grey bar in A). The white diamond indicates the average across subjects. **C)** The distributions show the difference between posterior distributions of stimulation conditions from the Bayesian mixed-effect model. A marginally credible difference was observed between posterior distributions after aMCC-TUS compared to PCC-TUS. **D)** The ERP curves show a late N2 component deflecting between 350 and 450 ms, which is highlighted by the light gray bar. The incongruent trials (full line) elicited more negative N2 relative to congruent trials (dashed line). The color pattern is the same as for the ERN. **E)** Betas of single-trial regression analysis were averaged in the above-mentioned time interval across participants for each stimulation condition. **F)** Comparison between posterior distributions showed credible difference between betas from sham relative to PCC-TUS and aMCC-TUS relative to PCC-TUS.

#### N2 and conflict monitoring was modulated by aMCC-TUS compared to PCC-TUS

As for ERN, the N2 component was present across all sessions (Fig. 6.D). Single-trial regression analysis estimates (Fig. 6.E) indicated that the congruence regressor had no credible effect in the sham condition (β = 0.14, 95% CrI [−0.32, 0.63]). However, relative to sham, aMCC-TUS was associated with marginally credible increased congruence betas (β = 0.10, 95% CrI [-0.02, 0.22]), whereas PCC-TUS showed credible reduced estimates (β = −0.12, 95% CrI [−0.20, −0.03]). Pairwise comparisons indicated that beta estimates (Fig. 6.F) were nearly credibly lower in the sham condition compared to aMCC-TUS (β = −0.101, 95% CrI [−0.22, 0.00]), and credibly higher compared for PCC-TUS (β = 0.116, 95% CrI [0.03, 0.2]). Importantly, beta values were also credibly higher in the aMCC-TUS compared to the PCC-TUS condition (β = 0.21, 95% CrI [0.09, 0.34]), suggesting that aMCC-TUS affected how response conflict modulates N2 amplitude relative to PCC-TUS.

#### aMCC-TUS modulated N2 and conflict monitoring relation over time

Next, we aimed to investigate whether TUS changed the N2-conflict monitoring effect over time. Note that this analysis was conducted only on the trials which survived the trial rejection of the Independent Component artefact correction. Similarly to the behavior results, we observed an interaction between Block and Stimulation (Fig. 7.A.1), with effect in the middle blocks relative to the early blocks after aMCC-TUS (β = 0.88, 95% CrI [0.02, 1.73]) compared to sham. However, the model also revealed a credible decrease in the estimate of aMCC-TUS relative to sham (β = −0.73, 95% CrI [-1.3, −0.16], Fig. A.2), confirmed by pairwise comparisons which showed a credible difference between sham and aMCC in early blocks (β = 0.72, 95% CrI [0.15, 1.3]). These findings are in line with the behavioural results which showed increased accuracy in early blocks, associated with a more negative conflict effect on the N2, and decreased accuracy in the middle blocks, associated with a more positive conflict effect on the N2 after aMCC-TUS. Therefore, these results showed that after 27 minutes of stimulation, the effect of TUS of aMCC changes in both behaviour and in electrophysiological markers like N2.

**Figure 7.**
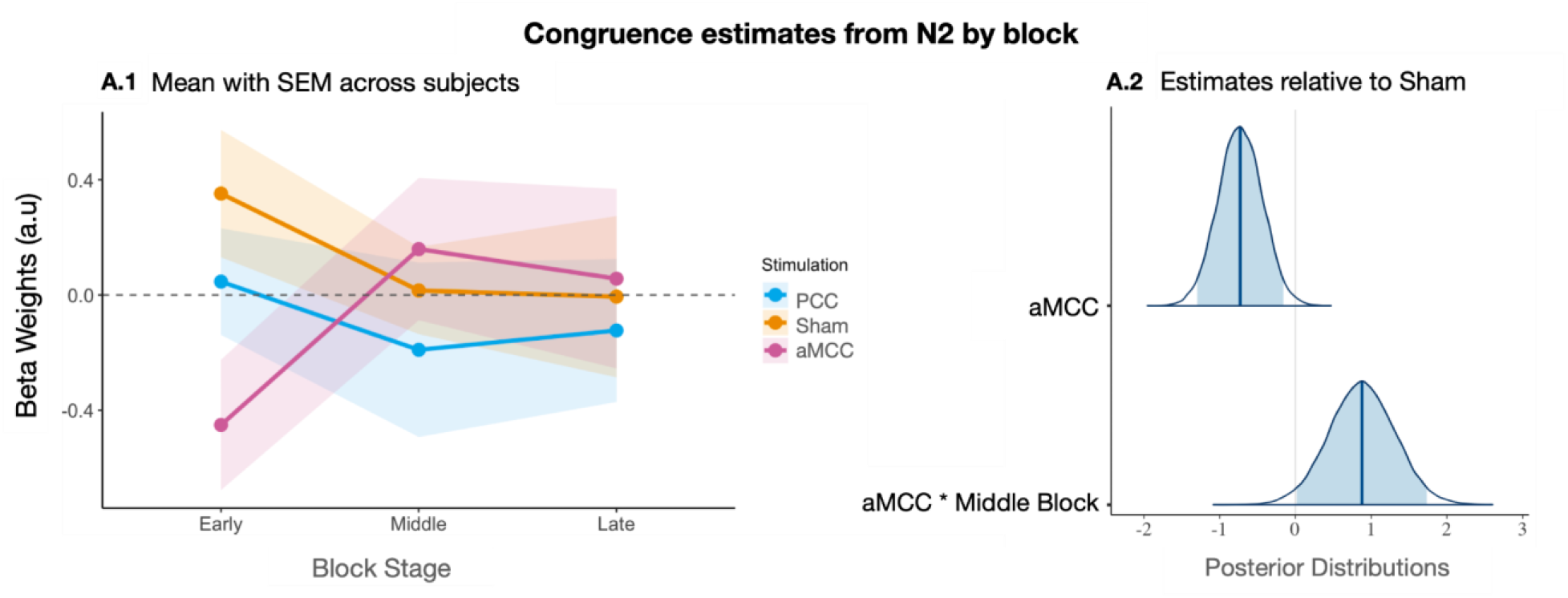
N2-Congruence relation by block: **A.1**) Beta weights of single trial regression analysis with grouped blocks showed different patterns after aMCC-TUS and sham condition, but not after PCC-TUS. **A.2**) Posterior distributions show smaller beta estimates after aMCC-TUS relative to sham, but larger betas in the middle blocks after aMCC-TUS.

### Drift Diffusion Model

In an exploratory analysis, we aimed to characterize the decision processes underlying the stimulation-dependent behavioral effects by fitting a multistage drift diffusion model (DDM) previously established and validated for this version of the flanker task. Details of the model have been described previously (Fischer et al., 2018; Kirschner et al., 2024) and are illustrated in Fig. 8.A. Model adequacy was evaluated using posterior predictive checks, which showed a good match between simulated and observed RTs and accuracies across multiple task factors (Fig. 8.C-F).

**Figure 8.**
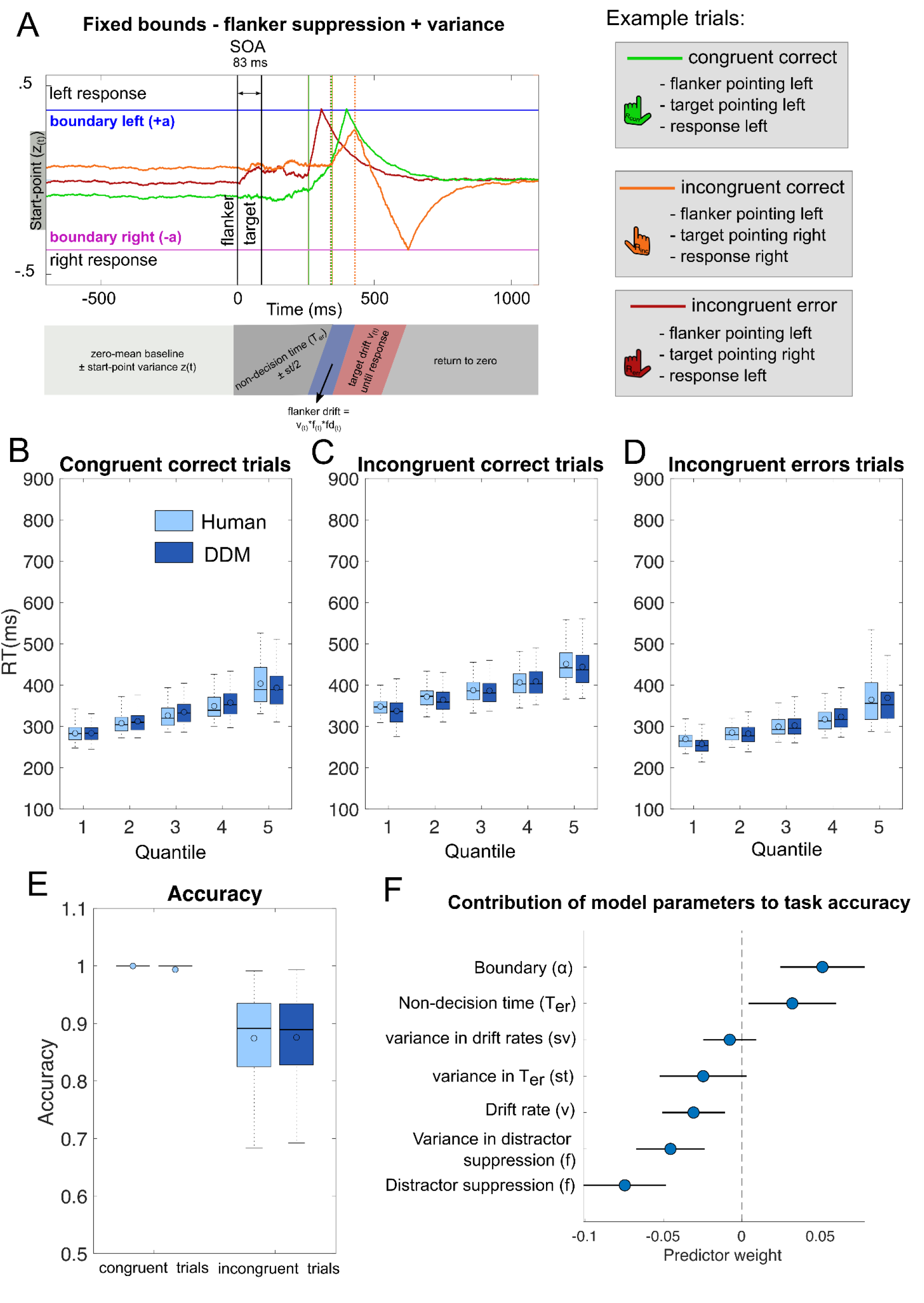
Multi-stage DDM captures task behavior and links model parameters to performance. **(A)** Description of the model: On a given trial, the decision can be randomly biased to favour one response (startpoint variance, sz_(t)_). The first stage of the model consisted of a pre-stimulus zero-mean baseline for which the value is determined by the start-point (z_(t)_) (Stage 1). After flanker presentation (0 ms), visual processing and motor execution times are captured by the non-decision time (T_er_) parameter, which can vary from trial to trial (st). In this period the decision process randomly drifts away from the start point (Stage 2). The next stage represents a noisy diffusion in flanker direction (f_(t)_) for the duration of the SOA which again can vary from trial to trial (sf) (Stage 3 - flanker diffusion). After flanker diffusion, stage 4 models the consecutive diffusion into the correct response direction with drift rate v_(t)_ with trial-to-trials variance (sv). Longer RTs for incongruent correct trials are due to an initial drift away from the correct response (orange decision process). Error likelihood increases when the baseline is shifted towards the flanker direction and f_(t)_ is higher on a given trial resulting in an early crossing of decision boundaries (red decision process). As a result, errors are usually faster on incongruent trials. Finally, Stage 5 models the return to baseline of the decision variable, akin to an Ornstein-Uhlenbeck process. The height of the boundary parameter (a) determines how much evidence accumulation is required to cross the boundary and trigger a response. (Figure adopted from Kirschner et al., 2024). **(B-E)** shows quantile fits of the model against human RT data (light blue) and model and human accuracy. In all conditions (congruent & incongruent correct as well as incongruent error), the model captures the RT data in each quantile, suggesting a good fit to the data. Note: boxes = interquartile range (IQR), o = median, - = mean, whiskers =1.5 × IQR, grey dots = outlier. **(F)** Regression coefficients and 95% confidence intervals (points and lines; sorted by value) stipulating the contribution of each model parameter estimate to overall participants task performance (i.e., overall accuracy on incongruent trials). The most important predictors were flanker suppression and decision boundary, with stronger suppression and a higher decision boundary associated with better performance.

To better understand the computational mechanisms modulated in our task, we first examined how task performance related to model parameter estimates. To this end, we regressed task performance onto an explanatory matrix containing parameter estimates across all conditions (Fig. 8.F). This analysis revealed that variability in several parameters was associated with overall task performance. The strongest predictors were distractor suppression and decision boundary. Here, stronger suppression—that is, lower (f) values, which reduce flanker-related drift (v_f_ = v × f × fᵈ)—and higher decision boundaries were associated with better performance.

Next, we investigated the effect of stimulation on parameter estimates across the whole task and during the early phase of the task (i.e., the first third, where we observed the strongest aMCC TUS effects). To facilitate convergence and improve parameter recovery, we fixed the variance parameters (sv, sz, st, sf) to the group mean. Parameter recovery analyses demonstrated that the fitting procedure reliably recovered the parameters used to generate synthetic data (see Supplementary Fig. 2).

The results of the parameter comparisons are shown in Fig. 9. Relative to sham, TUS of aMCC increased distractor suppression by ∼5% both across the entire task (0.409 ± 0.017 vs. 0.449 ± 0.016; increased suppression in 13/19 participants) and during the early task phase (0.409 ± 0.03 vs. 0.452 ± 0.027; 11/16 participants). Compared to PCC stimulation, aMCC stimulation was associated with slightly increased distractor suppression across the whole task (0.409 ± 0.017 vs. 0.427 ± 0.021; increased suppression in 10/19 participants) and early in the task (0.409 ± 0.030 vs. 0.444 ± 0.031; 10/16 participants). Three participants had to be excluded from the early-task analyses because they committed too few errors (<10) for stable model fitting. We also observed stimulation-related effects on decision boundary and drift rate, with lower drift rates and boundaries following aMCC stimulation relative to sham. However, because these parameters trade off against one another in the model, these effects are difficult to interpret mechanistically.

**Figure 9.**
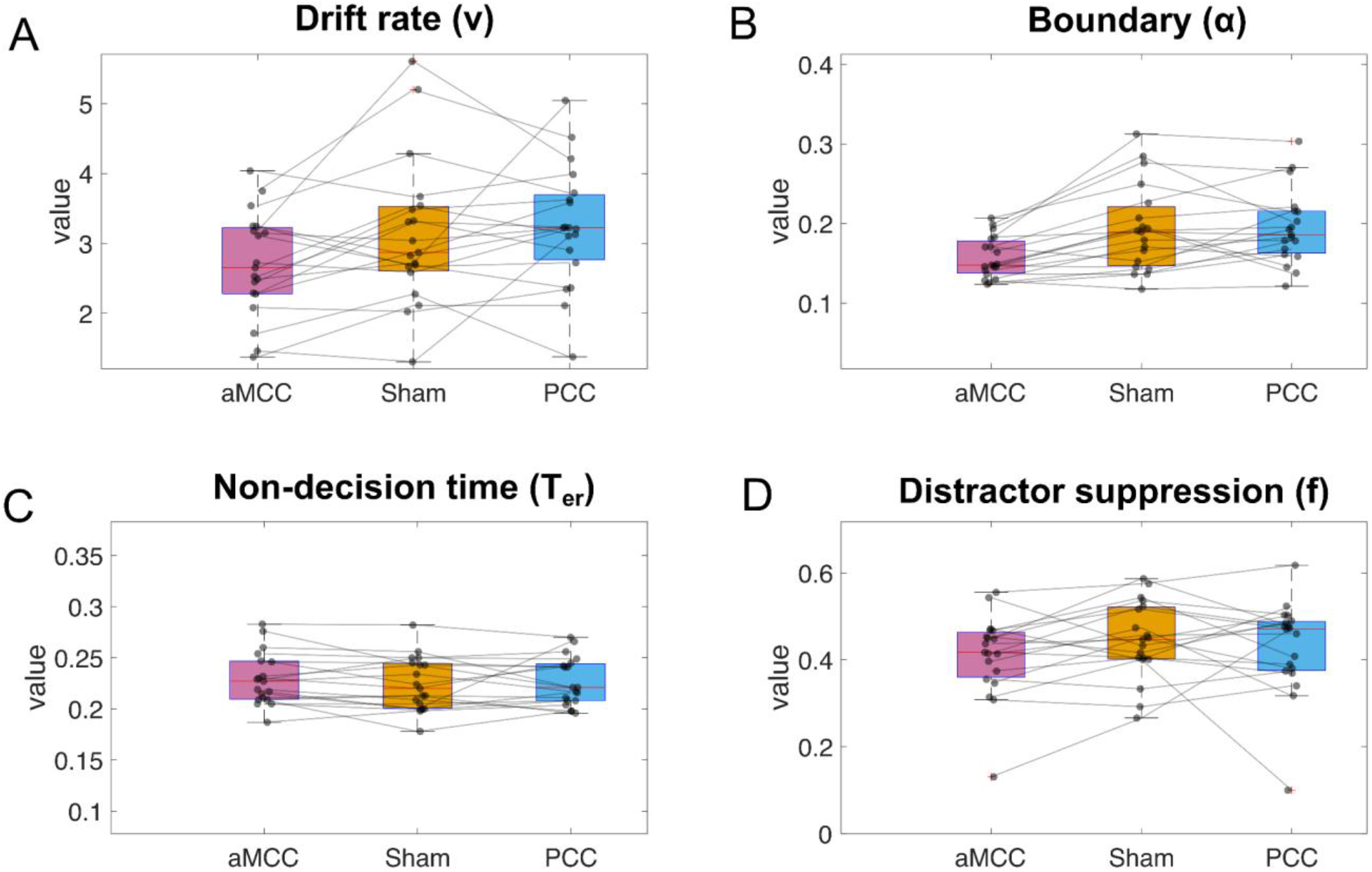
Parameter distribution across experimental conditions. **(A–D)** Comparison of parameter estimates from the winning model across stimulation conditions. **(A–B)** Stimulation-related effects were observed for decision boundary and drift rate, with lower drift rates and decision boundaries following aMCC stimulation relative to sham. However, because these parameters trade off against one another in the model, these effects are difficult to interpret mechanistically. **(C)** No stimulation-related effects were observed for non-decision time. **(D)** Relative to sham, TUS of aMCC increased distractor suppression by ∼5% and by ∼3% relative to TUS of PCC. Stronger suppression corresponds to lower (f) values, which reduce flanker-related drift (v_f_ = v × f × fᵈ).

Taken together, these findings complement the behavioral and neural results, suggesting that TUS of aMCC enhances accuracy by modulating conflict processing, as reflected in the effect on the N2. On a mechanistic level, the modeling results suggest that aMCC stimulation may enhance the suppression of distracting flanker information. However, these analyses were exploratory in nature and should be followed up in a larger sample. It should be noted that hierarchical mixed - effects analyses yielded relatively wide credible intervals that included zero, indicating that, with the current sample size, firm conclusions regarding the magnitude of stimulation effects on latent decision processes cannot yet be drawn.

## DISCUSSION

In this study, we investigated the causal role of aMCC in error processing and response conflict monitoring using 5Hz repetitive transcranial ultrasound stimulation. Several effects were observed which appeared to dissolve with increasing time past stimulation. TUS of aMCC increased accuracy while leaving reaction times unaffected. For PCC stimulation, a dose-dependent improvement in accuracy accompanied by a dose-dependent prolongation of reaction times was found. Furthermore, TUS of aMCC reduced the conflict effect on the N2. Contrary to our hypothesis, we did not find effects of TUS of aMCC (and PCC) on the ERN or on the difference between the waveforms for incorrect vs. correct responses. Notably, most of these results were present or strongest in the early time tercile of the experiment, reflecting the temporal dynamics of ultrasound effects on behavior and electrophysiological markers, consistent with findings previously reported in resting-state MRI studies. (Yaakub, et al., 2023; Atkinson-Clement et al., 2025b). By demonstrating an influence on electrophysiological markers and key cognitive control regions, such as the aMCC, these findings contribute to our understanding of the neural mechanisms underlying TUS and support further investigation of TUS as a potential intervention for disorders involving deficits in cognitive control.

### TUS of the aMCC increases proactive control and modulates response conflict

The frontocentral N2 has been shown to be modulated by the amount of response conflict (van Veen & Carter, 2002a; Folstein & van Petten, 2008; Danielmeier et al., 2009) and/or the control that is recruited upon detected conflict (Eichele et al., 2010). It has been suggested to be generated in the posterior medial frontal cortex, mainly the aMCC and the pre-SMA (Ullsperger & von Cramon, 2001, 2014a & 2014b; Wessel et al., 2012, van Veen & Carter, 2002b). The reduced conflict effect on the N2 and the increased accuracy suggests that TUS of the aMCC reduced either the amount of competition between the response tendencies induced by the distractors and the target, respectively, or the amount of control needed to resolve the conflict. The DDM analysis suggests that stimulation of the aMCC enhances distractor suppression, thereby reducing the influence of flanker-related information on evidence accumulation, which should result in reduced response conflict. Consequently, less reactive control would be needed upon incongruent trials. This would also mean that TUS of the aMCC increased proactive control thereby enabling more efficient suppression of the distracting input from the flankers. We can only speculate on the potential mechanisms of this enhanced distractor suppression. For example, stimulation of aMCC could potentially modulate attentional control, putatively via an effect of aMCC on cholinergic pathways from the basal forebrain to the visual cortices (Danielmeier et al., 2015, Ullsperger & Stork, 2021) or via direct connections from aMCC to the visual cortex, which have been demonstrated to play a role in attentional control in mice (Norman et al., 2021). Alternatively, changes in the excitation-inhibition balance between competing neuronal populations representing the distractor- and target-driven responses, respectively, could change the dynamics of choice selection (Wong & Wang, 2006). A shift of this balance towards excitation, demonstrated by increased glutamate and reduced GABA signals in the ventromedial prefrontal cortex, supported the convergence of the represented signal towards the chosen option in a value-based decision-making task (Jocham et al., 2012). It is tempting to speculate that our 5Hz rTUS protocol had a similar facilitatory effect increasing the excitability of the aMCC. A few previous reports support the notion of facilitatory stimulation effects on the aMCC. An early TMS study which targeted the aMCC during a Stroop task reported –similarly as in our study– a reduction in errors on incongruent trials, effectively abolishing the interference effect (Hayward, Goodwin & Harmer, 2004). However, it should be noted that TMS delivered to the more superficial left dorsal medial frontal cortex (SMA and pre-SMA) may disrupt performance by not only increasing reaction time but also error commission in incongruent trials during a Flanker Task (Taylor, Nobre and Rushworth, 2007). A recent TUS study using the same stimulation protocol showed increased connectivity of both, the aMCC and PCC after stimulation and a reduction of GABA in the PCC, suggesting a facilitatory effect (Yaakub et al., 2023).

Consistent evidence strongly suggests that the aMCC and pre-SMA are the main generators of the ERN (Debener et al., 2005; Fu et al., 2019, 2022; Godlove et al., 2011). In line with this, the ERN was absent in a patient with a focal lesion of the aMCC (Swick and Turken, 2002). Also, in patients with lesions of the ventrolateral anterior part of the thalamus, a region strongly projecting to the aMCC, the ERN is abolished (Seifert et al., 2011). However, to our surprise, TUS of the aMCC did not modulate the ERN. We speculate that the stimulation intensity may not have been sufficient to alter the neural activity patterns underlying the ERN. In other words, a stronger modulation of the aMCC may have been required to influence error processing itself, thereby producing changes in error-related activity following aMCC-TUS relative to the other conditions. In contrast to TMS, TUS does not induce depolarization of the targeted neurons. Instead, its effects may vary across brain regions due to differences in intrinsic connectivity, cytoarchitecture, and ion channel expression (Algermissen et al., 2026; Yaakub, et al., 2025; Farboud et al, 2026). Thus, with the low energy protocols that can be used safely in humans, no “virtual lesions” as in repetitive TMS can be induced. Such an effect might have been necessary to observe ERN modulation comparable to that reported in lesion studies. Furthermore, the subsequent Pe component was also unaffected by either aMCC-TUS or PCC-TUS relative to sham (data not shown), suggesting that the ERN–Pe complex could not be dissociated by our stimulation protocol. Perhaps an online TUS protocol would be more effective in directly influencing aMCC activity in the response window than an offline protocol and show direct effects on the ERN. Furthermore, our results indicate a dissociation of effects on the frontocentral N2 and the ERN, even though both are believed to be generated in the aMCC (Ullsperger et al., 2014a,b; Wessel et al., 2012; Gruendler et al., 2011). A similar dissociation was found in the patient with a focal aMCC lesion whose conflict-related N2 was unaffected while the ERN was abolished (Swick and Turken, 2002). This suggests that at least partly different neuronal populations seem to underlie these event-related potentials. It would be conceivable that TUS acts with different efficiency on different types of neurons. Moreover, it could be that the relative contributions of the aMCC and pre-SMA to the N2 and ERN, respectively, differ. Whereas the pre-SMA was spared in the lesion study, it was certainly co-stimulated in our TUS experiment.

### TUS of the PCC may lead to a dose-dependent speed-accuracy tradeoff

TUS targeted to the PCC also increased accuracy, even in a dose-dependent fashion, but, in contrast to the aMCC session, the conflict effect on the N2 remained unchanged. This suggests that the accuracy increase after PCC stimulation resulted from a different mechanism that left response conflict unaffected. Given the dose-dependency of reaction time prolongations across participants, a speed-accuracy trade-off appears to drive the effects of PCC stimulation. Although PCC is mainly associated with the default mode network, previous studies revealed that this region can actually be divided in at least two different parts with their specific roles (Foster et al., 2023; Leech et al., 2011, Leech et a.l, 2014). Foster and colleagues (2023), for instance, divided the PCC in ventral portion, which was found to be more integrated with the default mode network (DMN) and supports internally directed thought such as memory retrieval and self-referential processing; dorsal portion, which seems to be more engaged in executive and cognitive control processes, showing increased connectivity with frontoparietal (cognitive control) networks during demanding tasks; and retrosplenial cortex, which mainly involved with spatial memory and spatial cognition. Therefore, it is possible that our stimulation reached more strongly the dorsal PCC, hence the stronger modulation of behavioral responses. A recent meta-analysis revealed that the dorsal PCC is consistently activated during error processing (Cieslik et al., 2024). An MEG study (Agam et al., 2011) supports this interpretation by showing that the dorsal PCC, where the magnetic equivalent of the ERN was localized, may play a key role in detecting errors and generating the ERN, subsequently relaying this information to the aMCC to support adaptive behavioral adjustments. The authors reported that aMCC and PCC worked together on error processing as part of a coordinated functional network. Further, they demonstrated that functional connectivity analyses of these regions showed coupled activity both during task performance and at rest, consistent with their direct anatomical connections via the cingulum bundle. While TUS to the PCC did not reveal an effect on the ERN and Pe either, a credible difference between the effects of TUS to the PCC vs. the aMCC on the ERP difference between correct and incorrect responses was found, a finding that warrants further investigation. Taken together, our findings suggest dissociable contributions of these two regions to performance monitoring and cognitive control.

### Stimulation effects on neighboring regions

An important consideration about our findings is that we cannot rule out the effect of other regions potentially affected by the stimulation, due to the elongated cigar-shaped profile of the 250-kHz TUS beam. The stimulation may have also affected neighboring regions, such as the pre-SMA during aMCC stimulation or the precuneus during PCC stimulation. Nevertheless, we believe that potential co-stimulation of the pre-SMA does not fundamentally alter the interpretation of the aMCC findings, as both regions are thought to operate within a closely interconnected performance-monitoring network, as pre-SMA was revealed to play an important role in response inhibition and switching (Obeso et al., 2013), as well as in response conflict processing (Usami et al., 2013; Ullsperger & von Cramon, 2001). Supporting this view, a study in patients with intracranial implants identified subsets of neurons in both the aMCC and pre-SMA that signaled errors across different tasks, suggesting the existence of a domain-general error signal independent of sensory modality, motor demands, or conflict type (Fu et al., 2022). Importantly, the authors also reported temporal differences between these regions: response competition signals emerged earlier in the aMCC, whereas post-response signals appeared first in the pre-SMA and only later in the aMCC. In contrast, the effects of PCC-TUS on accuracy over time may indeed reflect modulation of the precuneus, a region strongly implicated in voluntary attentional shifts (Cavanna & Trimble, 2006).

In sum, our study provides evidence for a causal involvement of the aMCC in the allocation of proactive control that may support dealing with interference and response conflict. Moreover, it demonstrates that the 5Hz rTUS protocol may have facilitatory effects that show a specific temporal dynamics and decay within roughly 30 minutes. The ability to modulate recruitment of control may bear potential to influence maladaptive decoupling of heightened performance monitoring signals from appropriate control adjustments in mental conditions such as obsessive - compulsive disorder. The results furthermore point to a previously largely neglected role of the PCC in performance monitoring and cognitive control that is dissociable from the role of the aMCC. Future work will have to show whether different stimulation protocols targeting the aMCC can also affect error monitoring and post-error adaptations.

## DATA AVAILABILITY

Data will be made available upon request to C.S.A or M.U.

## ACKNOWLEGEMENTS

This project has received funding from the European Research Council (ERC) under the European Union’s Horizon 2020 research and innovation program (grant agreement No 101018805).

We would like to thank Denise Scheermann from the Neurology Department of Otto-von-Guericke University for the support with the MRI data acquisition.

## AUTHOR CONTRIBUTIONS

C.S.A. and M.U. conceptualized and designed the study. C.S.A. analyzed the fMRI, EEG, and behavioral data and performed the TUS simulations. L.V. contributed to the development of the TUS methodology, trained C.S.A. in TUS procedures, and provided laboratory resources for pilot testing. D.J. contributed to the behavioral modeling. H.K. conducted the drift diffusion modeling

(DDM) analyses. E.C. contributed to the development of the TUS simulation pipeline. C.S.A., M.U., H.K., and D.J. wrote the manuscript. L.V. critically revised the manuscript.

## SUPPLEMENTARY INFORMATION

S1. Priors based on the EEG session (no TUS)

S2. DDM parameter recovery.

## METHODS

### Participants

Nineteen participants (mean age=26 years, SD± 2.57, six women, one left handed man) with no previous history of psychological or neurological disorders volunteered to participate in a five-day study (one fMRI, three TUS+EEG and and one EEG-only session, whose results were used to estimate the priors for the behavior models). They were properly informed of the risks related to TUS application and signed a consent form approved by the Ethics Committee from Otto-von-Guericke Universität Magdeburg. Prior to every stimulation session, participants responded questionnaires to assess information such as hours of sleep and intake of illicit drugs. At the end of the session, they responded to a questionnaire to assess their state. Consequently, participants declared, on average, 7.01 hours of sleep (+-1.04), no medication or illicit drug consumption on the day of the experiment. All sessions, except the fMRI one, were kept in the same hour of the day within subjects.

### Sessions and Task

An arrow-based version of the Eriksen’s Flanker task, adapted from Fischer et al. (2018), was presented to participants across five sessions. During the non-fMRI^1^ sessions, participants were seated in a soundproof, dark room, positioned 70 cm from a screen. The task involved the presentation of four flanker arrows for 83 ms, followed by a central target arrow displayed for 33 ms. The orientation of the target arrow could be either congruent or incongruent with the flankers, and participants were instructed to respond as quickly as possible to the target’s orientation by pressing the “A” key with their left index finger or the “L” key with their right index finger. The arrows measured 0.46° × 0.8°, with 0.52° of space between them. Inter-trial intervals ranged from 0.75 to 1.5 seconds, following a Poisson distribution. The task included 720 pseudo-randomised trials, divided into 12 blocks (two participants performed 10/12 and two 9/12, due to technical problems), with half of the trials being congruent and the other half incongruent and lasted on average 34 minutes (SD=2.5 min).

### Functional Magnetic Resonance Imaging

Scanning sessions were conducted using a 3 Tesla Siemens PRISMA MR-system (Siemens, Erlangen, Germany) equipped with a 64-channel head coil. Functional data were collected from participants across 8 runs, comprising a total of 1448 volumes per subject. Blood oxygenation level-dependent (BOLD) signals were acquired with a multi-band accelerated T2*-weighted echo-planar imaging (EPI) sequence (multi-band acceleration factor of 2, repetition time (TR) = 2000 ms, echo time (TE) = 30 ms, flip angle = 80°, field of view (FoV) = 212 mm, voxel size = 2.2 × 2.2 × 2.2 mm, no gap, scan duration = 6.58 min). Volumes were acquired in an interleaved order, and consistent slice selection across sessions was ensured using the Head Scout Localizer, based on Autoalign (Siemens, Erlangen).

A high-resolution three-dimensional T1-weighted anatomical image was obtained using a magnetization-prepared rapid acquisition gradient echo (MPRAGE) sequence (T R = 2700 ms, TE = 2.42 ms, FoV = 230 mm, flip angle = 7°, voxel size = 0.9 × 0.9 × 0.9 mm, 225 slices, GRAPPA factor = 2, scan duration = 6.21 min). This anatomical map, covering the entire brain, served as a reference for registering the EPI data. Additionally, a pointwise-encoding time reduction with radial acquisition (PETRA) anatomical image (TR 1 = 3.32 ms, TE = 0.07 ms, flip angle = 1°, FoV = 294 mm, voxel size = 0.8 × 0.8 × 0.8, 353 slices, scan duration = 6.52 min) was acquired in order to subsequently use for the thermal and acoustic simulations.

Preprocessing steps counted with slice-timing, realignment (register to the mean), coregistration (to the T1) and smoothing (6 mm) and were conducted on SPM12 (www.fil.ion.ucl.ac.uk/spm, Wellcome Trust Centre for Neuroimaging, London, UK), as well as the GLM analysis. We used MarsBaR (Brett et al., 2011) to create the aMCC masks, which were a 3 mm sphere built based on the coordinates of the nearest local maximum inside the cluster resulting from the contrast of incongruent incorrect > incongruent correct trials, around the anterior portion of the MCC region. The PCC masks were similarly created, but the main voxel was selected based on the lack or lowest activity in the region.

### Transcranial Ultrasound Stimulation

The equipment used in this study was the NeuroFUS PRO system (Brainbox Ltd., Cardiff, UK) with a four-element ultrasound transducer (CTX-250-32, 64 mm diameter, Sonic Concepts Inc., Bothell, WA, USA) and a fundamental frequency of 250 kHz. We based our TUS protocol on the 5Hz protocol (Zeng et al., 2021, Yaakub et al., 2023). Specifically, we used a pulse length of 20 ms (plus 10 ms Tukey ramping), pulse repetition period of 200 ms, total duration of 120 s. The target ISPPA in water was kept constant at 30 W/cm2 for each participant, but the focus varied according to each subject’s anatomical structure (see Fig. 2.B1). Acoustic and thermal simulations were performed using K-Plan in order to assess the safety of stimulating the regions of interest in each participant before and after the stimulation session. For the simulation we used a T1w, masks of the aMCC and PCC and a pseudo CT image which was created based on an acquired PETRA image. For the pseudo CT transformation, we used SimNIBS alongside PRESTUS (https://github.com/Donders-Institute/PRESTUS, linear mapping option “carpino”).

Participants were invited to three TUS sessions, in which they received the 5Hz rTUS protocol for 120s targeting the aMCC or the PCC, here used as our active control region. Our sham stimulation consisted of a 2 second stimulation of the 5Hz-rTUS protocol targeting the aMCC. We prepared the participants’ hair with ultrasound gel, applying it layer by layer to ensure that no bubbles were trapped. The transducer was coupled using a water balloon containing distilled and degassed water, which was held in place by a plastic membrane and a 3D-printed ring (1 mm wide) supplied by Brainbox. After the stimulation was completed, participants had their hair washed and dried, an EEG cap was prepared with a reduced set-up (16 channels) before they started the flanker task. The time between the end of the stimulation and beginning of EEG measurement lasted on average 17 minutes (SD=1m50s) and the interval between stimulation sessions varied between 2 and 4 days. Table 1 depicts the average of the most relevant parameters from the post-session simulations according to (Martin et al., 2024). It is important to note that the stimulation parameters, such as Isppa and depth, were based on prior simulations, as those were tested and proved to be within the safe limits. However, the simulated depth did not always correspond to the exact depth required to reach our regions of interest, particularly due to the offset introduced by the a 3D-printed ring.

To address this issue, we created a ROI mask from the main cluster obtained from our contrast of interest (Fig. 2.A2). For each participant, the corresponding acoustic pressure map from each mask was obtained from K-Plan. To ensure spatial correspondence, the pressure volume was resliced into the native space of the ROI mask using trilinear interpolation via SPM’s reslice routine, and alignment was verified numerically before further analysis. Pressure values outside the ROI were excluded from all calculations. Within the ROI mask, the following metrics were extracted: peak pressure (maximum voxel value) and mean pressure.

A focal volume was defined by applying a −6 dB threshold to the full (unmasked) pressure map, that is, retaining all voxels whose pressure was at least 50% of the in-ROI peak. To isolate the sonication focus from spurious remote regions exceeding this threshold, a 26-connectivity component analysis was applied to the thresholded map, and only components with at least one voxel overlapping the ROI were retained. The intersection of this connected −6 dB mask with the ROI was then used to compute the mean pressure at the focus, the number of overlapping voxels, and the corresponding volume in mm³ (derived from the voxel determinant of the affine matrix). Full-width at half-maximum (FWHM) along each cardinal axis was estimated from the spatial extent of the −6 dB mask projected onto each dimension and converted to millimetres using the voxel resolution.

To guard against cases where the SPM-derived cluster fell entirely outside the intended hemisphere, for example due to a contralateral activation, an anatomical fallback was implemented. The aMCC and PCC masks were loaded for the relevant hemisphere. If no voxels overlapped between the SPM cluster and the hemisphere sphere, the sphere mask replaced the cluster as the working ROI. Acoustic pressure estimates extracted from the ROI analysis were subsequently used to derive standardised dose quantities. The absorbed dose was then calculated following Nandi et al. (2025). All dose quantities were computed in a fully vectorised pipeline applied to the subject-level dataset, with sonication duration assigned conditionally based on the stimulation region, thereby avoiding order-dependent indexing errors that could arise from sequential row-wise joins.

The values reported in Table 1 represent both the peak pressure within the intersection of the mask and the cluster, and the mean pressure within the −6 dB ultrasound beam. Our aim was to demonstrate that, even in the presence of a depth offset, the ultrasound beam still reached the main voxels within the mask and cluster with substantial intensity (Fig. 2.A1). We further calculated the absorbed dose (Fig. 2.A2), defined as Dose (or Exposure) = Isppa * PD, where Isppa is the spatial peak pulse average intensity, PD is the pulse duration (Nandi et al. 2025), to show that, although the simulations indicate that some intensity was delivered during the sham condition, the brief duration of two seconds was insufficient to result in a meaningful absorbed dose (left side = 1.0, ± 0.4; right side = 1.3, ± 0.56).

Finally, all individual masks were normalized to MNI space using ANTs software and plotted together (Fig. 2.A.4), so we could verify whether all participants received stimulation around the same portions of the anterior and posterior cingulate cortex.

### Electroencephalography

#### EEG acquisition and pre-processing

Preprocessing and first level EEG analysis were performed based on the analysis described in Fischer & Ullsperger, 2013; Fischer et al., 2018; Kirschner et al., 2022. Electroencephalographic signals were acquired using a reduced setup with 16 channels (FCz, FC4, C4, CP4, CP3, C3, FC3, AFC, PO4, PO3, LO2, T10, T9, LO1, IO1, IO2) from a montage with 61 Ag/AgCl sintered electrodes on five concentric rings equidistantly spaced around Cz. The vertical and horizontal central lines were identical to the positions of the 10% system. Ocular electrodes were placed below the left and right eyes and impedances were kept below 10 kΩ. The signal was continuously recorded at a sampling rate of 500 Hz BrainAmpMRplus amplifiers(Brain Products). Data were band-pass filtered offline between 0.3 and 20 Hz and segmented into epochs ranging from −1.5 to 2 s relative to target onset. The epochs were demeaned and analyzed using adaptive mixture independent component analysis (AMICA - Palmer, Kreutz-Delgado & Makeig, 2011). The time courses and topographies of the independent components of each dataset were visually inspected for components reflecting eye blinks, horizontal eye movements or electrode artifacts, and those components were removed from the data.

#### EEG analysis

For the statistical analysis, we conducted a single-trial regression analysis (with trials coded as: congruent = 1, incongruent = −1; correct response = 1, incorrect response = −1). To model the ERN, we selected only incongruent trials (both correct and incorrect) and included accuracy and RT as regressors. The accuracy regressor represents the relationship between EEG amplitude and the response; specifically, larger beta values indicate a greater amplitude difference between incorrect and correct responses. Note that the RT regressor is not presented in the results, as we did not have a specific hypothesis for it. To model the N2, we selected only correct responses from both congruent and incongruent trials, including congruence and RT as regressors. Similar to the ERN model, the congruence regressor indicates the relationship between signal amplitude and the conflict effect; larger beta values represent a stronger conflict effect in incongruent trials relative to congruent ones.

We employed Bayesian mixed-effects models (Eq. 1.1) to analyze the single-trial regression estimates of the ERN (β coefficients extracted from the 70–120 ms interval) and the N2 (β coefficients extracted from the 350–450 ms interval) at the FCz electrode. These intervals were selected based on the average peak activation for both ERPs for each stimulation condition individually. As exploratory analysis, we ran new single-trial regression analysis on grouped blocks (early = 1 to 4; middle = 5 to 8; and late = 9 to 12). We selected the betas from the same intervals as described above and fitted a new model, including stimulation and block stage as fixed and random effects (Eq.1.2). Note that for the four datasets which had fewer blocks, the grouping remained the same with the necessary adjustments.

### Behaviour analysis

Participants’ performance was assessed based on their reaction time (RT) and accuracy. All non- answered trials (113) and trials below 0.08s and above 0.9 s (37) were excluded from the analysis. In total 39511 trials were provided to trial-level linear and a generalised linear mixed models (binomial distribution with logit link) in which log-transformed RT and Response (correct vs. Incorrect), respectively, were predicted. Stimulation (verum [aMCC], sham [aMCC], active [PCC]) and congruence (congruent vs. incongruent) conditions were used as fixed effects and subject as random effect. We also included block, trial number and previous trial in different models, but since they did not yield conclusive results, we refrained from including them here. Behaviour analysis was conducted in R-Studio using brms (Bürkner, 2017.) and emmeans library (R version 4.4.0 - https://www.r-project.org/). Informative priors were defined based on the data of non-TUS- EEG sessions (Supplementary Table 1).

In order to investigate the effect of stimulation on accuracy and reaction time, we adopted the models described below (Eq. 2.1 and 2.2) and controlled for additional factors. Subject-specific random intercepts and random slopes for stimulation were included in the random-effects structure, allowing the different effects to vary across participants to account for individual differences in behavioral sensitivity to stimulation. Both models showed sufficient convergence (all R^ = 1.00). In further analysis, we explored different time windows of the TUS effect by dividing the task in three block stages (early = 1-4; middle = 5-8; late = 9-12) by fitting Eq. 2.3. All two-level variables were coded as −0.5 and 0.5 for the estimates to directly reflect the difference between them. We used the sham stimulation as reference in the three-level stimulation parameter, hence all the estimates should be interpreted accordingly.

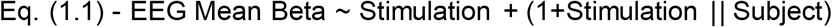

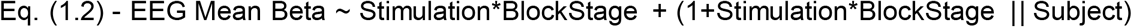

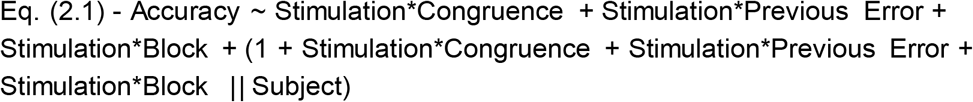

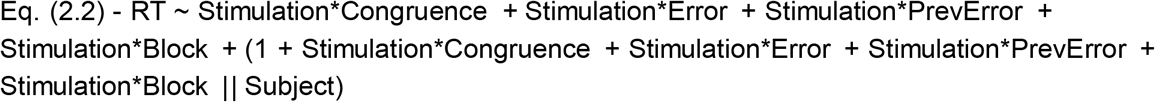

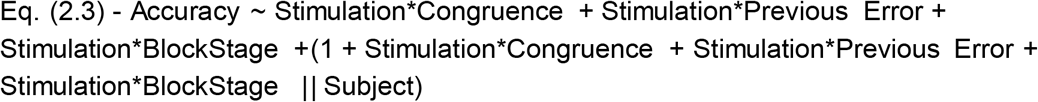

### Correla1tions

The ISPPA used in the correlations were calculated based on the ROI peak pressure (Table 1), as we aimed for the direct correspondence between the focus of the stimulation, the delivered intensity and the behavioral outcome, thus potential noise coming from voxels outside our regions of interest. All correlations were carried out using R-project software.

### Drift–diffusion modelling

Basic features of the multistage sequential sampling model and general fitting procedures were described by Fischer et al., (2018) and Kirschner et al., (2024). The description is adapted therefrom.

#### DDM stages

Our DDM assumed that the decision signal in the flanker task undergoes several stages (Fig. 8A). The start point of each trial was modelled as a baseline period prior to flanker onset (Stage 1). Here, each time step of the decision signal was drawn from a normal distribution with a mean of zero and an SD of 0.001 and shifted by a model-free variance parameter (sz) reflecting start points at each trial. This period was followed by a second stage during which the decision signal could drift randomly away from the start point for the duration of the nondecision time (Ter) of each trial (Stage 2). We allowed for noise accumulation during this period because we previously demonstrated that quick random diffusion in the nondecision time partly explains simple response errors in response conflict tasks (Fischer et al., 2018). Thereafter, we assumed that evidence accumulation was driven by flanker direction for as long as these were displayed (Stage 3). Consecutive to the flanker diffusion, evidence accumulation was driven by the direction of the target stimulus until the response threshold was met (Stage 4). Finally, we modeled a consecutive return of the decision signal to zero according to an Ornstein–Uhlenbeck process (Uhlenbeck & Ornstein, 1930) (Stage 5). In short, the Ornstein–Uhlenbeck process is a stochastic process that reverts a signal to a mean, θ, with speed, κ, and volatility, σ. We fixed these parameters to θ=0, κ = 0.004, and σ = 0.001. To speed up the model fitting procedure, we neither simulated baseline periods nor returned to zero during the fitting because these stages have no effect on model predictions and were merely added to the model to facilitate comparison between the predicted decision signal from the DDM to the time course of the EEG signal (see Fischer et al., 2018 and Kirschner et al., 2024).

#### Model parameters and description

As in most DDMs, evidence accumulation in our model is governed by a Wiener process with stepwise increments according to a Gaussian distribution with mean v (called drift rate) and within-trial variance s (reflecting the system’s noise, i.e., the amount of noise per computation step of the diffusion), which was fixed to 0.1. The step size for all models was set to 1 ms. A decision (response) is triggered when the diffusion reaches a criterion (Fig. 8A threshold or boundary, blue and magenta lines, respectively, with values ± α). We used symmetrical boundaries that were defined as left-hand responses when the positive boundary was reached first and as right-hand responses when the negative boundary was reached first (Fig. 8A). While only flankers were on screen, the model’s diffusion was driven by the direction of the flanking arrows (i.e., positive when left, negative when right), scaled by a free parameter f. Specifically, the drift rate during the flanker-only period (v1) on a given trial t was determined by the following:

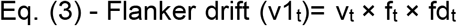

Here, v_t_ and f_t_ on a given trial t were drawn from a normal distribution with mean v and f and their associated variances (sv and sf, respectively, see below); fd_t_ reflects the direction (+1, flanker pointing to the left; −1, flanker pointing to the right) of the flankers on a given trial t. After the target onset, the diffusion was governed by the target direction. This drift rate (v2, whereby v2_t_ = v_t_) reflects the combined influence of the target plus flanker on the decision signal. Moreover, we assumed that despite the disappearance of target and flanker arrows after 33 ms of common presentation, the decision continues to form with a constant speed (v_t_). This was based on several studies that indicate that constant drift rates account well even for cases in which visual input is masked after a certain period of time (Ratcliff & Rouder, 2000). Therefore, left-pointing incongruent flankers (Fig. 8.A orange drift lines) are associated with a positive drift during the SOA, but a negative drift thereafter. Note that both periods are shifted in time by the nondecision time modeled as a free parameter (Ter). Ter here simulates the translation time of stimulus evidence into decision formation, which can be expected to mainly reflect visual processing and response mapping, and varies to a certain degree on every trial, for example, due to fluctuations of alertness. Specifically, across trials, we assume that the start point (z) of evidence accumulation (the baseline), nondecision time (Ter), drift rate (v), and distractor suppression (f) vary to some degree. As the task comprised exactly 50% left and right responses, we fixed the mean start point (z) of the diffusion process to 0. The model was thus unbiased regarding the average starting point across the experiment. However, each individual trial’s starting point was allowed to randomly vary according to a uniform distribution with lower and upper limits fit as the free parameter sz x 0.5 (i.e., upper/lower limit = 0 ± sz/2). This variance in start points can be interpreted as the range of bias of a participant toward a left- or right-hand response that varies between trials. Moreover, we modeled the nondecision time (Ter) single-trial variance with parameter st as a uniform distribution with borders = Ter ± st/0.5. This simply reflects that in some trials, evidence may take longer to be processed, for example, via fluctuations in attention. To account for variance in the drift rate, we allowed drift rates to vary according to a zero-mean Gaussian distribution with variance sv. Those trials with higher drift rates will reach a decision quicker yet may also be more prone to reach the incorrect boundary on incongruent trials, where the flanking stimuli point away from the correct direction. Additionally, we assumed that the degree to which distractors influence the diffusion process may vary between trials, reflecting selective attention and attention slips. Therefore, we modeled trial-by-trial flanker suppression effects according to a zero-mean Gaussian distribution with variance sf. In sum, our model comprised four free parameters (v, f, a, Ter) and four trial-by-trial variance parameters (sv, sf, sz, st).

#### Model fitting

We estimated the parameters for our model by fitting its parameters to RT and accuracy data observed using quantile maximum likelihood statistics (Heathcote et al., 2004) and differential evolution algorithms (Price et al., 2005). All congruent error trials were omitted for model fitting, as is common practice when fitting sequential sampling models to RT (Vandekerckhove & Tuerlinckx, 2007). Specifically, for each subject, we split the RTs into 10 equal-sized quantiles to estimate quantile maximum likelihood statistics (which is similar to a χ^2^ statistic) minimizing the negative log-likelihood of the observed participant data given each set of model parameters. The likelihood of each single observed RT is determined by the model’s likelihood of predicting an observation in the corresponding bin separated by correct and erroneous responses. Additionally, we used a mixture model assuming 2% contaminates that were distributed uniformly over the full range of RTs in correct and error responses. For model fitting in all iterations, we applied the following hard priors, which can be seen as boundary parameters: v (0.01–8.5), sv (0–1.5), a (0.01–0.45), sz (0.05–0.3), Ter (0.1–0.4), st (0–2), f (0.1– 1.5), and sf (0–1.5).

## SUPPLEMENTARY MATERIAL

S.1. We opted for not including the non-TUS EEG session in our main models because of the difference in the procedure compared to the TUS sessions (e.g. no hair preparation, neuronavigation, etc.). However, we fitted a bayesian mixed-effects model to the data in order to determine the priors for the reaction time and accuracy models. Therefore, the obtained priors are depicted in table 1.

|  | Acc (sd) - Bernoulli | RT (sd) - Lognormal |
| --- | --- | --- |
| **Intercept** | normal(3.14, 0.6) | normal(-1.119, 0.2) |
| **Congruence** | normal(1.55, 0.6) | normal(-0.1450, 0.2) |
| **Error** | - | normal(0.2004, 0.2) |
| **Previous Error** | normal(0.037, 0.6) | normal(-0.0015, 0.2) |
| **Block** | normal(0.0144, 0.6) | normal(0.0069, 0.2) |
| **Trial** | normal(0.068, 0.6) | normal(0.0105, 0.2) |
| **Feedback** | normal(0.019, 0.6) | normal(0.0016, 0.2) |
| **Stimulations** | normal(0, 0.6) | normal(0, 0.2) |
| **Interactions** | normal(0, 0.6) | normal(0, 0.2) |

Baseline included in the model:

Model results revealed a robust effect of congruence (β = 2.24, 95% CrI [1.86, 2.61]), indicating higher accuracy for congruent relative to incongruent trials. No credible main effects were observed for stimulation, previous error, or block progression, as all corresponding credible intervals overlapped zero.

Among the interaction terms, only the interaction between Congruence and Stimulation2 reached credibility (β = 0.40, 95% CrI [0.01, 0.80]), suggesting that the congruency effect was enhanced under this stimulation condition relative to the reference condition. No credible interactions were observed between stimulation and previous error or between stimulation and block progression, although the interaction between Stimulation3 and block progression showed a trend towards decreasing performance across blocks (β = −0.14, 95% CrI [−0.29, 0.01]). Overall, these findings indicate that behavioural performance was primarily driven by conflict processing, with limited evidence for stimulation-related modulation.

### S2. Parameter Recovery

**Supplementary Figure 1.**
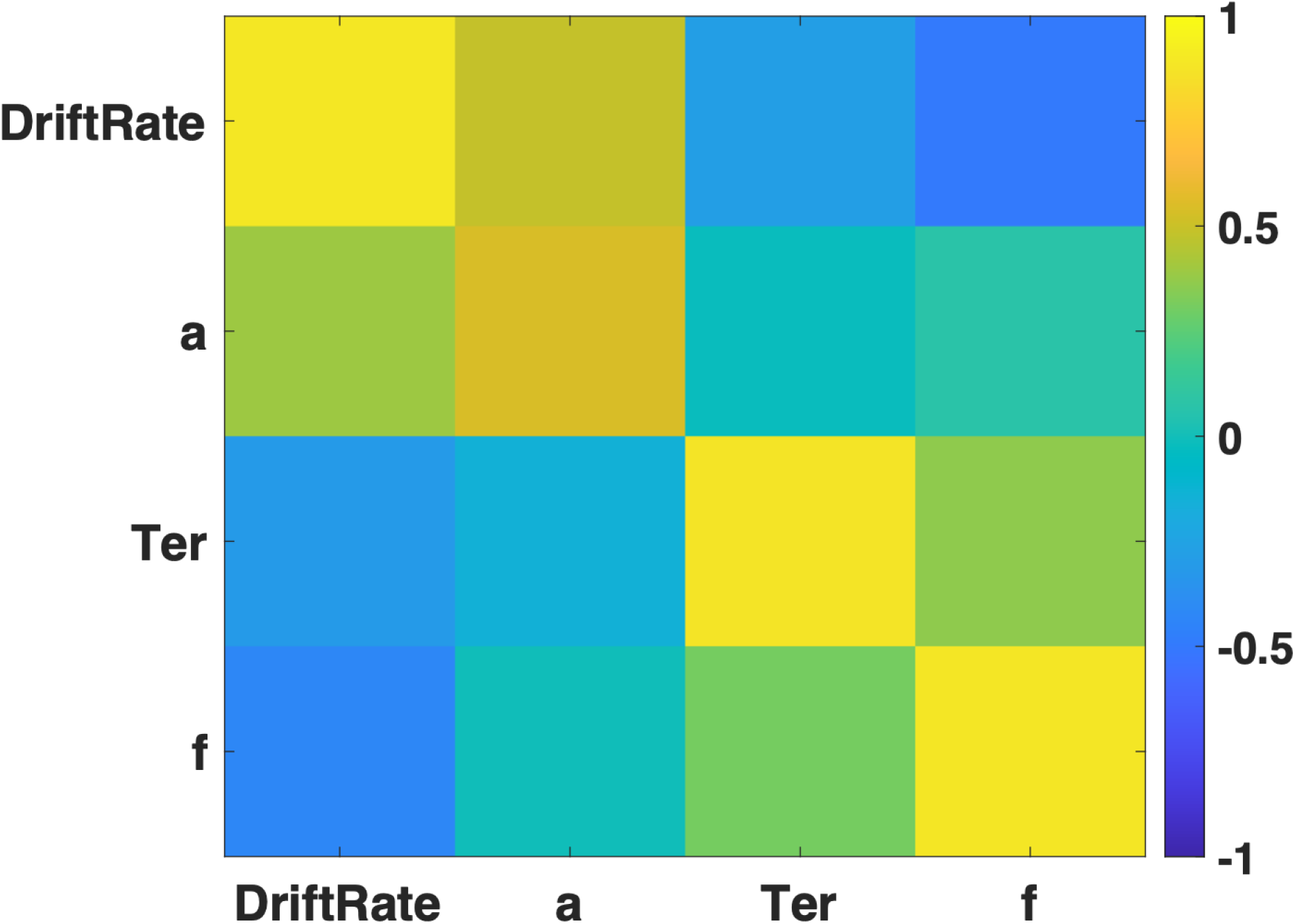
Parameter recovery. For parameter recovery analyses, we randomly drew model parameters out of a Gaussian distribution with mean and variance equal to the observed fitted parameters across the whole group to reduce parameter value combinations that were extremely unlikely to occur in human data. We simulated 1.000 parameter combinations and used the same differential evolution algorithm to recover the fitted models. Models that produced no errors at all or for which constraints were not met, were discarded from analysis. As in the human data, we used 5.000 trials per simulation. Plotted are correlations between simulated and recovered parameters for. Results indicate that parameter values that were used to simulate data from the full feature DDM (ordinate) tended to correlate with the parameter values best fit to those synthetic datasets (abscissa).

## Footnotes

1 The task was adjusted for the fMRI session. Participants were positioned 35 cm from the screen, resulting in the arrows appearing slightly larger. Inter-trial intervals (ITIs) ranged from 4 to 6 seconds, following a Poisson distribution, and an additional 16 seconds were added at the end of each run to account for the decay of the BOLD signal. A total of 480 trials were divided into 8 blocks. The number of trials was reduced to accommodate the increased ITI duration, ensuring the experiment remained a manageable length.

